# FMRI multi-scale cortical spontaneous activity: 7T vs. 3T

**DOI:** 10.1101/2021.06.09.447694

**Authors:** Xiu-Xia Xing, Chao Jiang, Xiao Gao, Yin-Shan Wang, Xi-Nian Zuo

## Abstract

This paper describes the use of the Human Connectome Project (HCP) data for mapping the distribution of spontaneous activity in the human brain across different spatial scales, magnets and individuals. Specifically, the resting-state functional MRI signals acquired under the HCP 3 tesla (T) and 7T magnet protocols were measured by computational methods at multiple spatial scales across the cerebral cortex using: 1) an amplitude metric on a single measuring unit (ALFF), 2) a functional homogeneity metric on a set of neighboring measuring units (ReHo) and 3) a homotopic functional connectivity metric on pairs of symmetric measuring units between the two hemispheres (VMHC). Statistical assessments on these measurements revealed that all the raw metrics were enhanced by the higher magnetic field, highlighting their dependence on magnet field strength. Measurement reliability of these global measurements were moderate to high and comparable between between 3T and 7T magnets. The differences in these measurements introduced by the higher magnetic field were spatially dependent and varied according to specific cortical regions. Specifically, the spatial contrasts of ALFF were enhanced by the 7T magnet within the anterior cortex while weakened in the posterior cortex. This is opposite for ReHo and VMHC. This scale-dependent phenomena also held true for measurement reliabilities, which were enhanced by the 7T magnet for ReHo and VMHC and weakened for ALFF. These reliability differences were primarily located in high-order associate cortex, reflecting the corresponding changes of individual differences: higher between-subject variability and lower within-subject variability for ReHo and VMHC, lower between-subject variability and higher within-subject variability for ReHo and VMHC with respect to higher magnetic field strength. Our work, for the first time, demonstrates the spatial-scale dependence of spontaneous cortical activity measurements in the human brain and their test-retest reliability across different magnet strengths, and discussed about the statistical implications for experimental design using resting-state fMRI.

## Introduction

Mapping the intrinsic architecture of human brain function in vivo can be accomplished using functional magnetic resonance imaging (fMRI) to estimate the spontaneous brain activity at a resting-state (rfMRI) [6, 7, 17, 35]. The growing field of network neuroscience can use the fMRI derived data to model the entire set of functional interactions in the form of the human connectome [3, 42, 46]. This functional connectomics has greatly enhanced our knowledge and understanding of brain organization (e.g., a small-word, rich-club, modular and gradual connectome [8, 9, 25, 47]) and how connections are optimized for mind and behavior (e.g., intelligence [2]). These general observations have largely been made at the group level, and challenges are emerging when translating observations into individual-level functional connectomics [27, 41], particularly when quantifying an individual’s differences in their functional connectome [14, 18].

In measuring individual differences, two concepts are fundamental, namely reliability and validity [61]. Reliability characterises a proportion of measurement variability between different individuals from the overall variability including both between- and within-individual (a.k.a random) variability components [51]. It is commonly used to reflect the consistency or agreement of the measurements among different occasions. However, it can also be a measure reflecting the discriminability. In other words, for a measure to sufficiently capture (i.e., discriminate between) individual characteristics, the reliability will need to be high. Discriminability is dampened in measures that underestimate between-individual variability, even when the consistency of the measurements within-individuals are equal. Thus a higher reliability is essential for the measurement to better differentiate a group of individuals, i.e., inter-individual differences [61]. Recent work has proven that measurement reliability is equivalent to the fingerprint and the ability of the measures to distinguish individuals under the Gaussian distribution [31]. It also provides an upper bound of the measurement validity [4], which cannot be as readily quantified as the reliability [61]. High-level reliability is thus the primary requirement for measuring individual differences, the quantification of which is important for guiding individual-level research [33, 61] into developmental trends [24] and clinical variations [30].

Previous meta-analyses have demonstrated that test-retest reliability can vary substantially among different rfMRI measurements [32, 59]. These differences reflect the complexity in handling various individual variability measurements related to the three aspects of functional connectomics: biological target, imaging tool and computational metric. Specifically, high-order associative target (e.g., networks) of human brain function are more variable across different subjects (higher between-individual variability) and thus more reliable. Structural targets (e.g., cortical thickness) are more reliable than functional targets (e.g., functional connectivity) due to their lower within-subject variability (i.e., more stable across different measurement occasions within individuals). We also observed that some strategies on imaging tools and computational approaches such as scanning duration and global signal regression impact the rfMRI measurements and their reliability [60]. Based on previous findings [59, 61], we can make three major observations: 1) metrics with a structural or morphological basis are more reliable than those without; 2) local metrics are more reliable than global metrics; and 3) advances in rfMRI acquisition protocols (spatial-temporal resolution and scan duration) improve test-retest reliability. We recently demonstrated the benefits of the latter (#3) using data from the Human Connectome Project (HCP) [48], to determine increased short-term (two-day) test-retest reliability with the advanced protocol [59]. The previous reliability assessments on rfMRI measurements of the human brain were all derived from 3T magnets. MRI acquisition protocols are continuing to advance, and many brain imaging studies are now being conducted using 7 tesla high-field strength magnetic fields. It is not yet known how the improvements in data quality (i.e., the signal-to-noise ratio) from 7T acquisitions [48, 49] affect the test-retest reliability of rfMRI measurements, and thus our ability to study individual-level differences in human cortical spontaneous activity (CSA).

Here, we employed the test-retest HCP rfMRI datasets from the same group of healthy adults scanned twice at both 3T and 7T HCP magnets. The aim of our analyses is to compare differences in common CSA metrics of the human connectome between the HCP 3T and 7T rfMRI settings in terms of their regional variations, individual variability and test-retest reliability. Specifically, we chose three CSA metrics at multiple spatial scales across the cerebral cortex: 1) amplitude metric on a single measuring unit (ALFF) [54, 55], 2) functional homogeneity metric on a set of neighboring measuring units (ReHo) [53, 60] and 3) homotopic functional connectivity metric on pairs of symmetric measuring units between the two hemispheres (VMHC) [58]. These CSA metrics are determined as the present research of interests according to their high reliability and potential validity [26, 59]. We first perform a replication of the previous work on test-retest reliability using the HCP 3T data [59], and then update the evaluation based on the HCP 7T data and finally, test the differences between the HCP 3T and 7T magnets.

## Materials and Methods

### Participants and Test-Retest Data

Among the 1200 participants of HCP 1200 release, a total of 84 participants (age range: 22-36 yrs; mean age: 29.6 yrs; 32 males) received 4 rfMRI scans during two days at the 3T and 7T HCP scanners, respectively. A final list of all the participants is available at Github (ADD LINK). Each participant received a pair of high-resolution (0.7mm isotropic voxel) structural images of T1-weighted (T1w) and T2-weighted (T2w) sequences only at the WU-Minn Connectome 3T scanner. One-hour rfMRI signals were acquired with a two-day test-retest design to measure the SCAs. Specifically, this rfMRI protocol comprises of four 15min-scans (two scans per day). Table 1 demonstrated details of the rfMRI scanning sequences across the 3T Connectome scanner and the 7T MAGNETOM scanner. More details can be found in two recent papers [48, 49].

### Data Preprocessing

All preprocessing steps for both 3T and 7T images were implemented using the HCP pipeline, which generated all the publicly shared images via the HCP database. These steps have been comprehensively described in previous HCP publications [19, 20, 40], and we only briefly highlight several key steps here.

Temporal filtering was applied with a minimal high-pass filter of a 2000s cutoff, which is almost equivalent to removing a linear trend from the data. An ICA-based approach, FIX, was used to remove non-neural spatio-temporal artifacts from each 15-min rfMRI scan. Performance offered by FIX has been fully described and evaluated in previous publications [36]. Specifically, ICA was first carried out using MELODIC to decompose the volumetric rfMRI data into a set of components [5, 57]. FIX classifies these components into signal and noise components with a training-validation scheme, and further cleanup the rfMRI scans by regressing out the noise components from the data. All the FIX processing steps are implemented in the volumetric space, and the resulted data are then projected into the MNI152 space with the HCP gray-ordinate system. Spatial smoothing was performed with a small Gaussian kernel of 2mm on the surface to avoid the tissue-mixture effect on both signal and noise within the volume space while preserving the boundaries across different functional units.

In the present work, we constrain the data analyses to the cerebral cortex [13, 60]. Specifically, 3T rfMRI data are projected onto a left-right symmetric cortical surface grid, which is reconstructed with the HCP-customized FreeSurfer pipeline based upon 69 healthy adults and comprises 32,492 vertices per hemisphere with an approximate 2mm inter-vertex distance, namely Conte69_LR32k. For 7T rfMRI data, an averaged surface models was derived from the 62 HCP participants to establish a left-right symmetric cortical surface model with a higher spatial resolution (~ 59k vertices per hemisphere), namely HCP62_59k. The preprocessed rfMRI data were then projected onto the HCP62_59k grid. To make direct comparisons of the rfMRI metrics between 3T and 7T magnets, we down-sampled all individual 7T rfMRI surface data onto the Conte69_LR32k surface grid for the subsequent rfMRI computation of the three CSA metrics. A group-level surface mask was established by including every vertex showing rfMRI signals of both 3T and 7T datasets from all the four rfMRI scans of the 84 participants across the cortical mantle.

### Raw Metric Computation and Normalization

Vertex-wise metrics allow for direct and high-resolution characterization of the intrinsic functional architecture of the human cerebral cortex with respect to its organizational morphology and function. Three widely used metrics (ALFF, ReHo and VMHC) are employed to characterize the CSA in the human brain across different spatial scales. These metrics have been defined and described in detail in our recent work on test-retest reliability evaluation [12, 59].

The three metrics were calculated based on the slow-4 band (0.033Hz - 0.083Hz) CSAs of each voxel in the group-level mask [10, 21]. The amplitude of low frequency fluctuation (ALFF) of the CSA at a single voxel was derived (~ within 2mm distance) [54]. The regional homogeneity (ReHo) characterizes the local functional connectivity across the cortical mantle [26, 60]. To quantify the ReHo vertex-wise, the Kendall’s coefficients of concordance (KCCs) of the rfMRI time series among the 4-step neighboring (~ 61 vertices within 16mm distance) vertices were calculated.Voxel-mirrored homotopic functional connectivity (VMHC) was derived as the temporal correlation (Fisher-z transformed) between the rfMRI timeseries from a pair of symmetric voxels between the two hemispheres (> 16mm distance) [58]. As in Figure 1, these metrics measure CSAs across different spatial scales (i.e., from regional, local activity to distributional, distant connectivity) [37].

**Figure 1.**
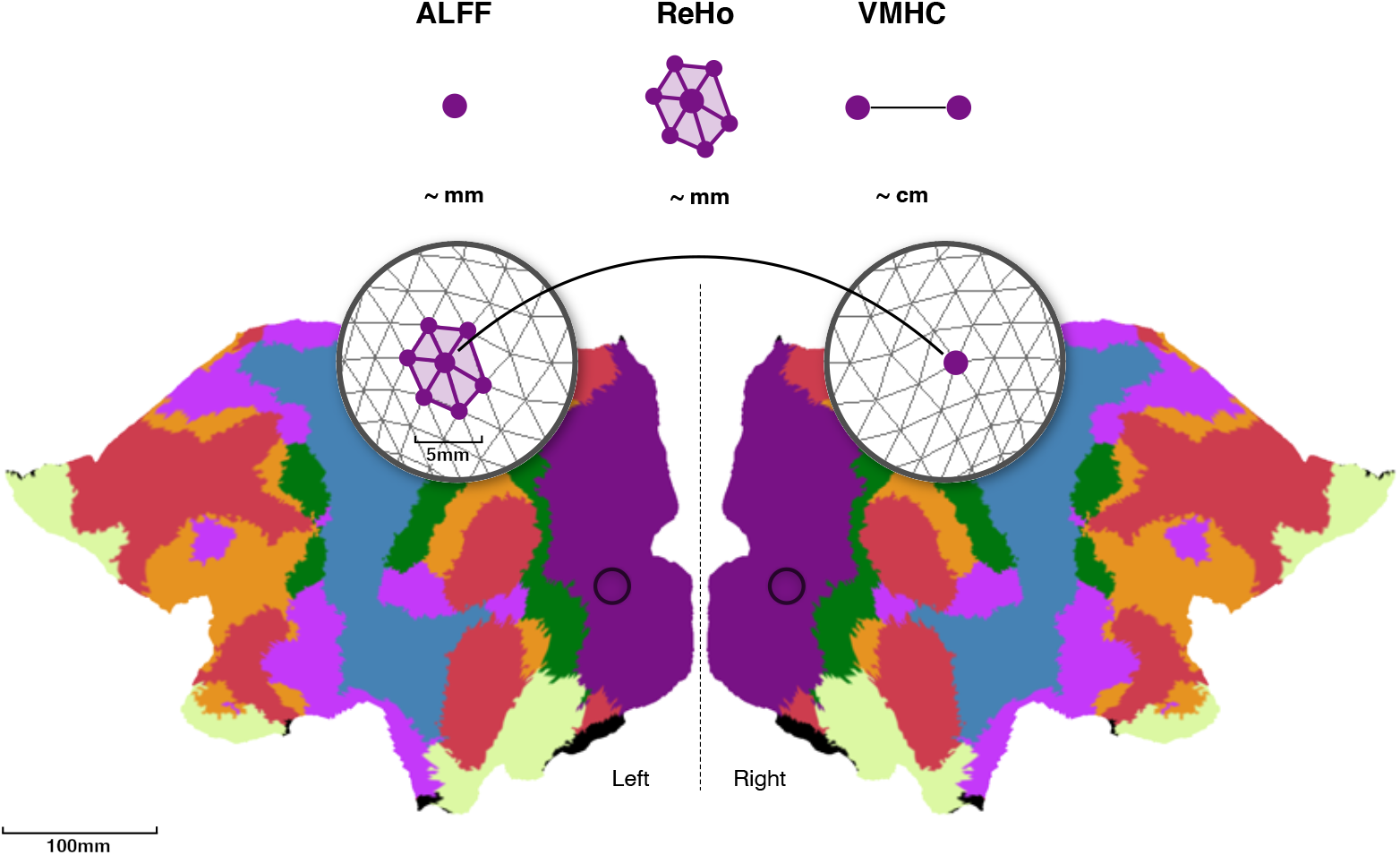
Multiscale CSA metrics. ALFF, ReHo and VMHC are calculated for each vertex at the cortical surface of each hemisphere. ALFF measures CSA at single vertex (within a 2mm location). ReHo measures local connectivity of CSA among a set of neighboring vertices (within a 16mm area). VMHC measures distant connectivity between the symmetrical hemispheric pair of vertices (outside of 16mm distance). The Conte69_LR32k surfaces are left-right symmetrical and visualized with grids for the two hemispheres where the colors indicate the seven intrinsic connectivity networks [44].

For each rfMRI scan, both the mean and standard deviation (SD) of the raw maps for each metric are first calculated within the group mask. Individual raw metric maps are then converted into z-value maps by subtracting the mean and dividing by SD. The z-value map can be considered as a gradient (relative value) map, normalized to the global mean and SD. This normalization allows these maps to be comparable across scans and field strengths.

### Group Patterns

To examine the degree of differences induced by the 7T magnet compared to the 3T magnet, we performed a set of paired t-tests on the rfMRI-derived CSA metrics described above. Specifically, the individual mean and SD metrics of the two scans (taken in one day) were averaged to obtain a single set of individual means and SDs for the metrics to increase the signal-to-noise ratio and reduce the complexity of the comparisons. The average operation was also applied to the two individual z-value maps obtained from scans taken the same day. A paired t-test on each global metric was performed between 3T and 7T magnetic field strengths to test if the overall CSA metrics are different between the two scanner field strengths. Similar tests were applied to the individual z-value maps at the vertex level. Constrained within the group mask, vertex-wise paired t-tests compared all averaged individual maps between 3T and 7T, and produced the general statistics. A quantity of the test significance sign(*t*) × (– log_10_(*p*_corrected_)) was used to visualize the degree of strength on differences in various metrics between 3T and 7T. The final significance maps were corrected for multiple comparisons with the Bonferroni method (uncorrected *p* < 0.05/*N*, where *N* is the number of vertices within the mask).

### Individual Differences

Linear mixed effect (LME) models were used to estimate both intra-individual and inter-individual variability of CSA metrics at global (mean and SD) and voxel (z-value map) level. One strength of the linear mixed models is capable of explicitly separating the age or sex related variability with the repeated CSA measurements by including them as covariates. We modeled both age and sex as participant-level covariates due to the very short test-retest duration (two days). We averaged the metrics derived from the two 15-min rfMRI scans within the same day for each participant to feed in the LME models described as in the equation (1).

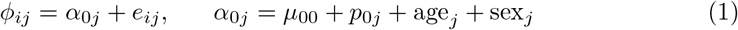

This equation denotes *ϕ_ij_* as the i-th day measurement of the *j*-th participant (*i* = 1, 2; *j* = 1,…, 84) where *μ*_00_ is a fixed parameter of modeling the group mean measurement across all the participants’ repeated measurements. The term *p*_0*j*_ models the participant effect whereas sex_*j*_ and age_*j*_ can monitor gender and age effect, respectively. The term *e_ij_* is the random error. These models were implemented and solved in R with the **lme** package. Specifically, the variance of *e_ij_* was estimated as the within-participant or intra-individual variability (*V*_W_) while the variance of *p*_0*j*_ was estimated as the between-participant or inter-individual variability (*V*_b_). As in Figure 2, to offer an intuitive way of understanding the changes of individual variability and reliability from 3T to 7T, we introduce and update a reliability-plane visualization we introduced previously [51]. The ICC values are categorized into five common intervals: 0 ≤ *R*_ICC_ ≤ 0.2 (slight), 0.2 < *R*_ICC_ ≤ 0.4 (fair), 0.4 < *R*_ICC_ ≤ 0.6 (moderate), 0.6 < *R*_ICC_ ≤ 0.8 (substantial), 0.8 < *R*_ICC_ ≤ 1 (almost perfect).

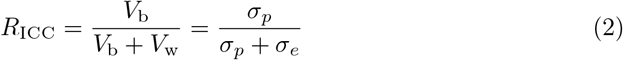

**Figure 2.**
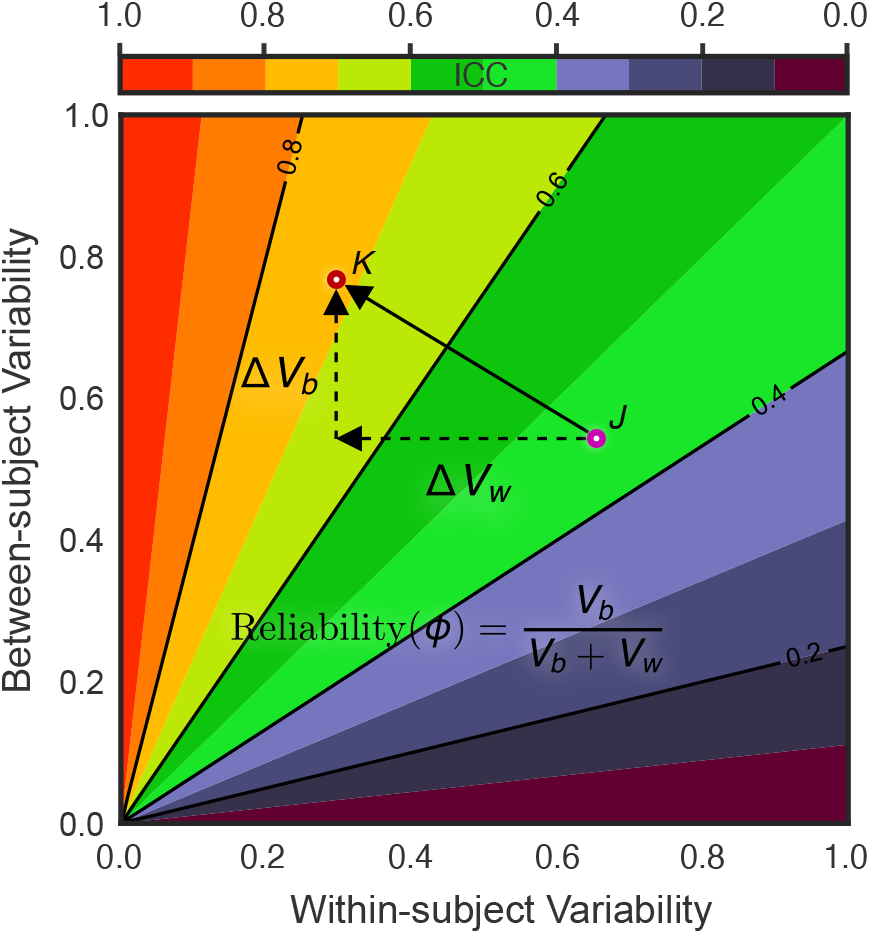
Reliability gradient as a function of individual variability changes. Both *V*_b_ and *V*_w_ are normalized by the total sample variances to have values between 0 and 1. Their changes (Δ*V*_b_ and Δ*V*_w_) introduce a reliability gradient as represented by the vector (black arrow). The length of the arrow reflects the amplitude of reliability differences when the reliability assessment changes from one choice (pink circle, *J*) to another choice (red circle, *K*). The arrow’s direction (**JK**) indicates the sources of this reliability change. Here the reliability goes from moderate to substantial as the Δ*V*_b_ increases and the Δ*V*_b_ decreases.

These variance components were estimated by implementing the restricted maximum likelihood (ReML) approach to avoid negative values of the intra-class correlation (ICC), which is defined as in the equation (2). All these models were equally performed to the three rfMRI-derived CSA metrics from both 3T and 7T HCP datasets.

## Results

Our analyses generated many results, and we will present these findings according to group patterns (spatial distribution) and individual differences (reliability assessment) in terms their differences between 3T and 7T magnet.

### Group Patterns of Cortical Spontaneous Activity

Paired t-tests revealed that the HCP scans taken using a 7T magnet produced globally higher CSA metrics (i.e., mean ALFF, ReHo and VMHC) than those taken using 3T magnets across the three different spatial scales (Figure 3). The spatial variability of these metrics as measured by their SD across the cortical mantle increased from the 3T magnet to the 7T magnet, but not for the ALFF metric. These findings are reproducible across the two days.

**Figure 3.**
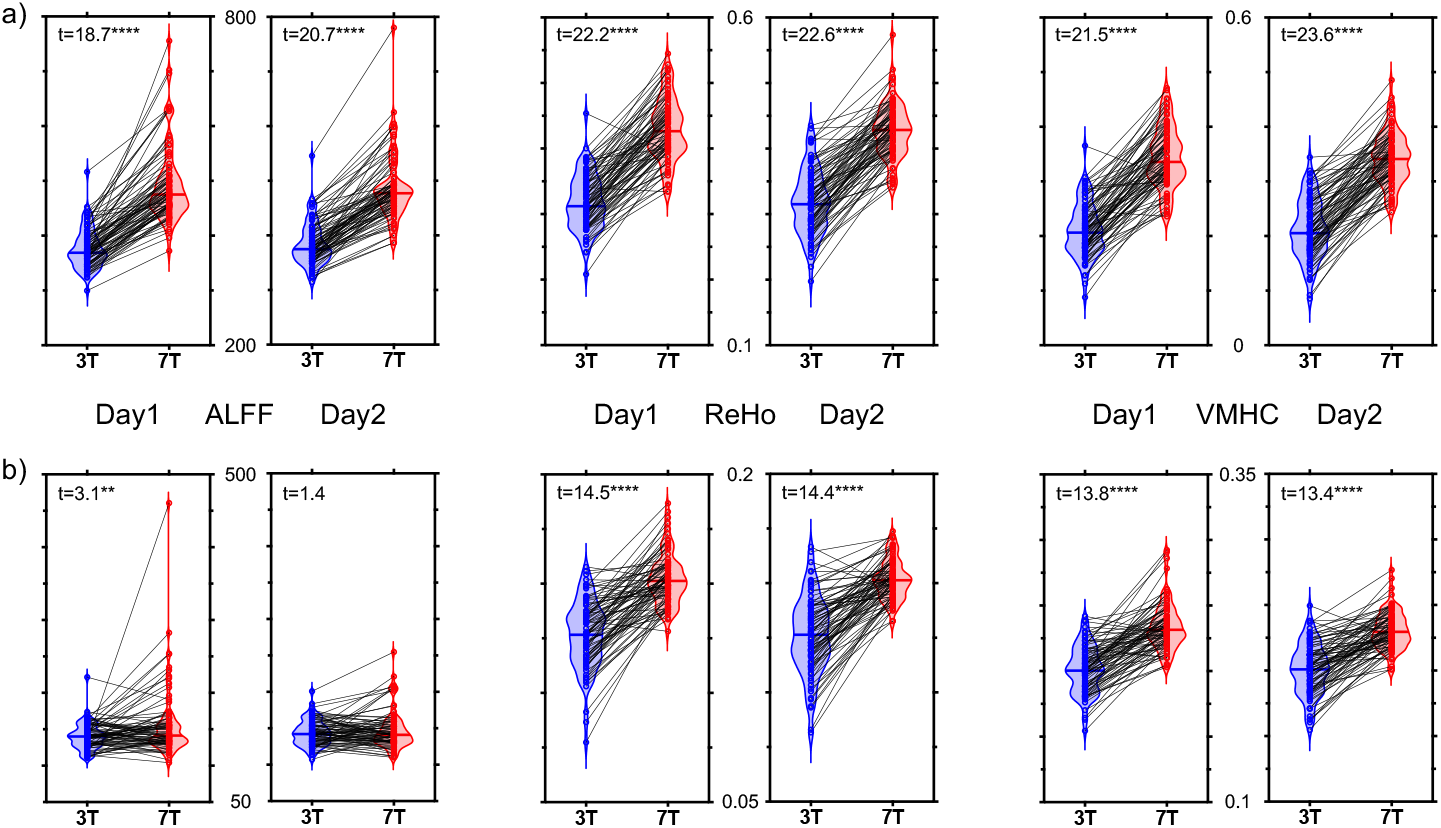
Paired plots of multiscale CSA metrics between 3T and 7T magnet. Violin plots are presented for global mean (a) and SD (b) of ALFF, ReHo and VMHC across Day1 and Day2 in the HCP-style imaging protocol. The t-statistics for each paired t-test is labeled in the left-up corner of the corresponding box while the related significance is shown as ** (*p* < 0.01), *** (*p* < 0.001) or ***** (*p* < 0.0001).

These signal enhancements at the global level by the 7T introduced different changes of the regional gradient level or rank (i.e., the z-value) between amplitude and connectivity metrics (Figure 4). Vertex-wise mean z-values of the three CSA metrics are rendered in Figure S1 for the 3T and 7T scans respectively. ALFF exhibited significant field strength-related changes of z-values along a spatial distribution from posterior to anterior cortex while the occipital and parietal areas showed increases of their z-values but temporal, frontal and insular areas demonstrated deceases of their z-values (Figure 4a, left). The lateral parietal cortex received the most significant z-value increases whereas the insula, medial prefrontal cortex and temporal pole were with the most significant z-value decreases. At network-level, the most spatial locations (more than 20% vertices) with increased z-values are distributed in dorsal attention and sensory motor network but default, limbic and ventral attention network for decreased z-values (Figure 4b, left). In contrast, ReHo (Figure 4a, middle) and VMHC (Figure 4a, right) distributed their z-value changes by the 7T magnet from anterior to posterior in the lateral cortex but from dorsal to ventral in the medial cortex. For ReHo, most increases happened to the default and ventral attention network while most decreases sit in the limbic and visual and default network (Figure 4b, middle). For VMHC, z-values increased in the ventral attention, sensory motor and default network but decreased in the limbic, default and visual network (Figure 4b, right). These findings are also provided under a 400-parcels of the network parcellation of the human cortex (see Figure S2).

**Figure 4.**
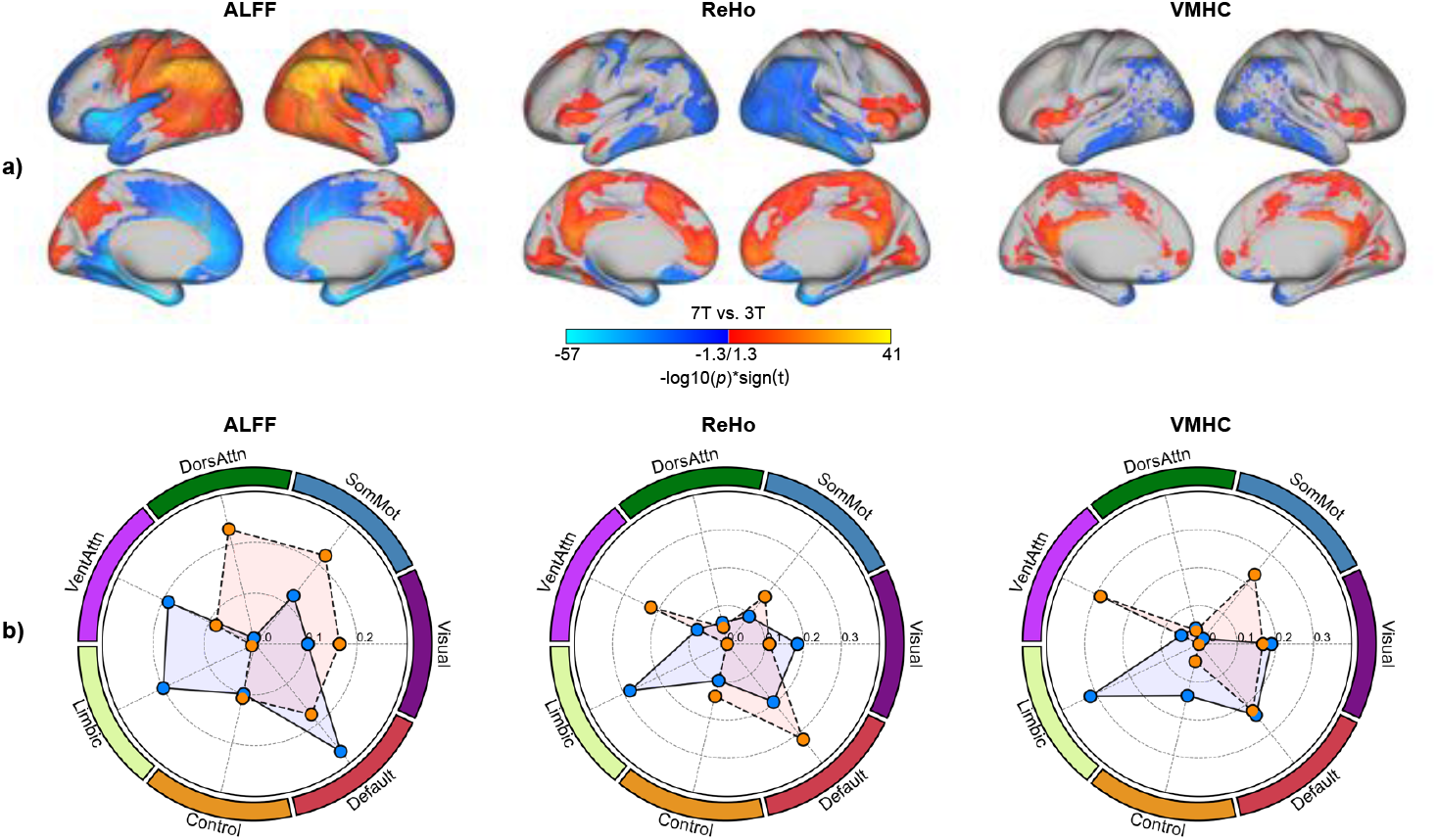
Surface and network mapping of differences in multiscale CSA metrics’ z-value between 3T and 7T magnets. a) Statistical maps of the paired t-tests on the three metrics are rendered onto the four cortical surfaces (the lateral and medial surfaces in both left and right hemisphere). The color indicates the significance on the tests (warm color: 7T higher than 3T; cool color: 7T lower than 3T). b) Proportion of significant vertices are summarized according to the seven canonical large-scale brain networks: Visual (visual), SomMot (sensory motor), VentAttn (ventral attention), DorsAttn (dorsal attention), Control (frontal parietal control network), Limbic (limbic) and Default (default). More details of these networks can be found in [44].

### Individual Differences in Cortical Spontaneous Activity

The changes in individual differences in global CSA metrics (mean and SD) from the 3T to the 7T magnet were generally subtle (Figure 5). Specifically, the variability of mean ALFF between subjects was smaller with the 7T magnet than with the 3T magnet, while the corresponding SD variability between subjects was almost identical between the 7T and 3T magnets. In contrast, for large-scale connectivity metrics including ReHo and VMHC, the between-subject variability of their mean metrics slightly increased from the 3T to 7T magnets, while the within-subject variability decreased. The direction of variability changes for the connectivity measurements (ReHo and VMHC) was opposite that of the ALFF. Regarding the test-retest reliability, these global CSA metrics all demonstrated moderate (0.4 ≤ *R*_ICC_ ≤ 0.6) to substantial (0.6 ≤ *R*_ICC_ ≤ 0.8) reliability. We noted that the mean ALFF exhibited an ICC of nearly 0.8, which indicates close to an almost perfect reliability. Reliability changes were limited to 0.1 of the ICCs for all the global metrics (except for the SD of ALFF) from the 3T to the 7T magnet.

**Figure 5.**
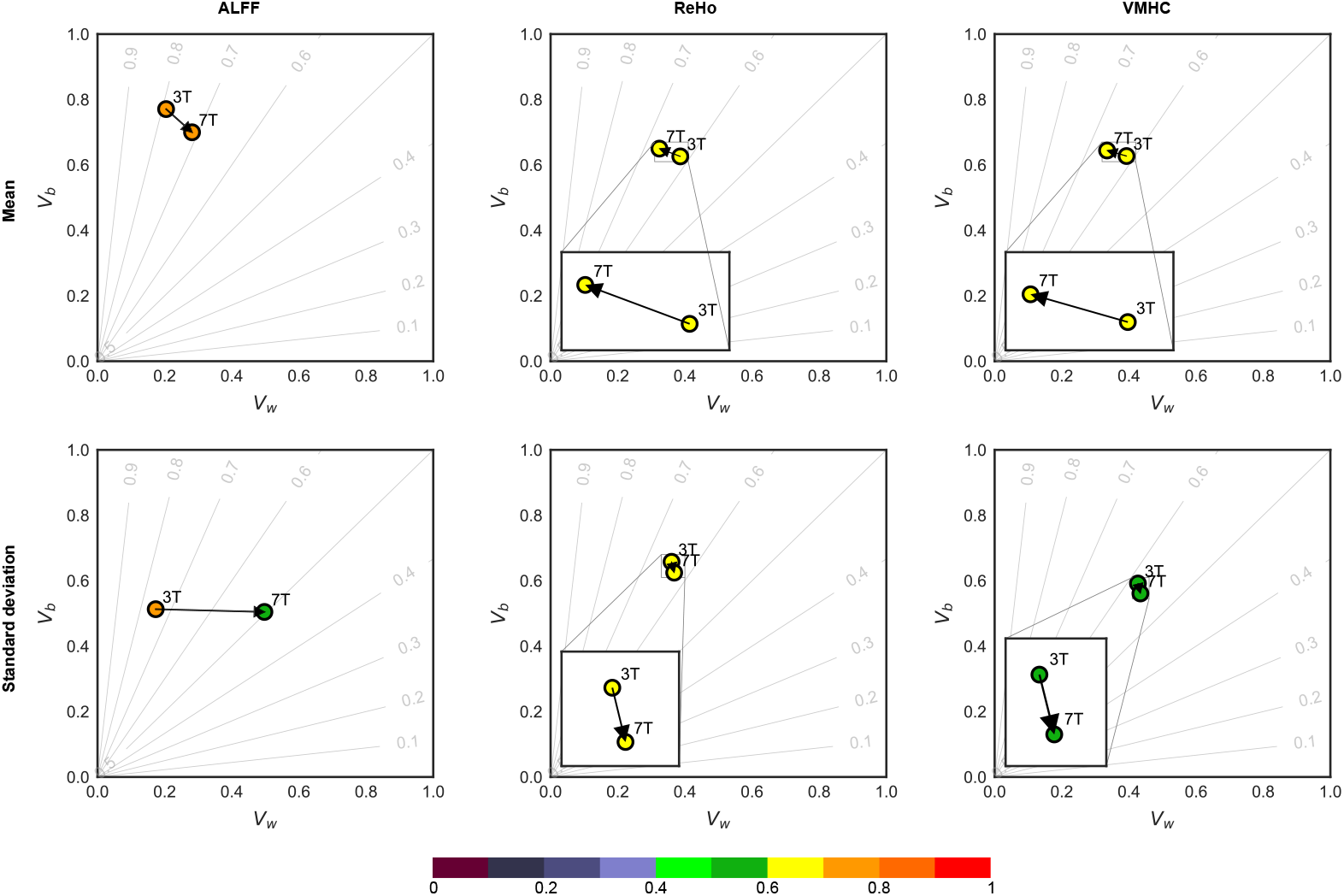
Reliability gradient of multi-scale CSA metrics from 3T to 7T magnet. Measurement reliability are plotted as small circles as function of both between-subject variability *V_b_* and within-subject variability *V*_W_. The values of ICC are outlined with an interval of 0.1 in the plane and assigned the corresponding colors in the colorbar. The face colors of these circles indicate their ICC levels.

Regarding the test-retest reliability, the 7T magnet (Figure 6b) reduced the reliability of ALFF and increased the reliability of both ReHo and VMHC compared to the 3T magnet (Figure 6a). In particular, VMHC gained the most observable improvements in its measurement reliability under the 7T magnet. At the network-level (Figure 6c), the largest changes in ALFF reliability were in the dorsal attention, control, default and sensory motor networks. In contrast, the most evident increases in ICCs for ReHo and VMHC were in the limbic, dorsal attention, default, control and sensory motor networks. We noted that such reliability improvements happened in the VMHC of the ventral attention network and not in the ReHo.

**Figure 6.**
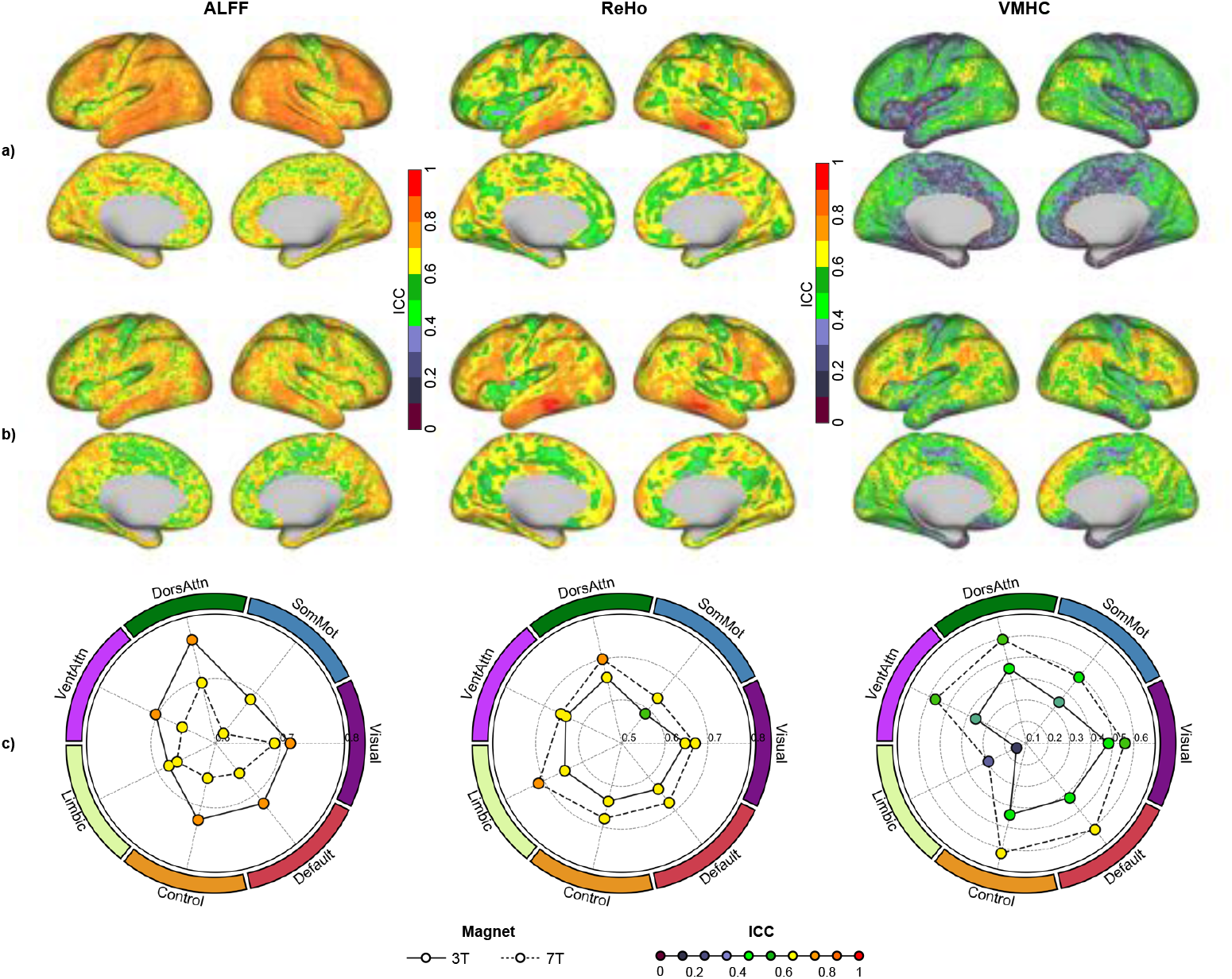
Surface and network mapping on test-retest reliability of multiscale CSA metrics under 3T and 7T magnet. a) Reliability maps of the three metrics derived with the 3T HCP magnet are rendered onto the four cortical surfaces (the lateral and medial surfaces in both left and right hemisphere). The color indicates the level of the ICCs. b) Reliability maps of the three metrics derived with the 7T HCP magnet are rendered onto the four cortical surfaces. c) Mean ICCs of all vertices within the seven canonical large-scale brain networks are plotted for both 3T (solid line) and 7T (dash line) magnet while the face colors of these circles indicate the level of ICC: Visual (visual), SomMot (sensory motor), VentAttn (ventral attention), DorsAttn (dorsal attention), Control (frontal parietal control network), Limbic (limbic) and Default (default) [44].

These patterns of reliability changes can be attributed to individual differences in z-values (Figure 7). Such patterns are highly similar to those observed at the global level (Figure 5) but provide more details on the differences in their spatial distribution (Figure S3). Specifically, the 7T magnet increased the within-subject variability and decreased the between-subject variability of ALFF z-values, but reversed for both ReHo and VMHC z-values (Figure 7a). At the network level, across both the 3T and 7T magnets, high-order associative networks, including the control, default and dorsal attention networks, were more variable among subjects than those primary networks (Figure 7b) and more stable within subjects (Figure 7c). From the 3T to the 7T magnets, the sensory motor network showed remarkable within-subject variability (Figure 7b, left) and the reduced between-subject variability (Figure 7c, left) in ALFF z-values. The ReHo of the limbic network exhibited large improvements in the between-subject variability (Figure 7b, middle) and reduced the within-subject variability of ReHo z-values under the 7T magnet (Figure 7c, middle). The ventral attention network was more recognizable among subjects in terms of the VMHC z-values under the 7T magnet (Figure 7b, right) and more stable across the two days within individual subjects (Figure 7c, right). These findings were also provided under a 400-parcels of the network parcellation of the human cortex (see Figure S4).

**Figure 7.**
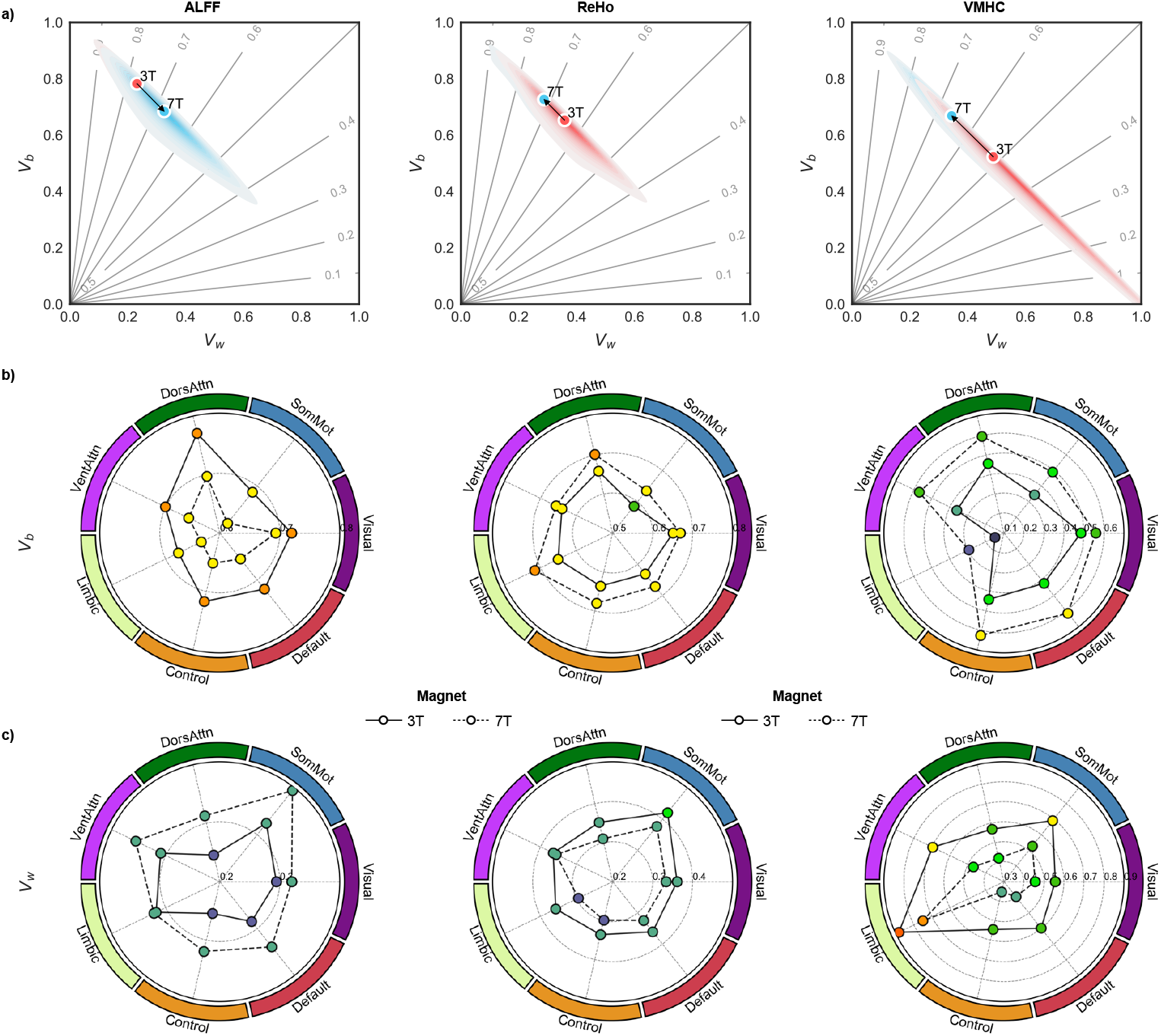
Reliability plane and network mapping on individual variability of multiscale CSA metrics from 3T to 7T magnet. a) Reliability gradient of multiscale CSA metrics from 3T to 7T magnet. Measurement reliability of each vertex is plotted as one member of the cloud as function of both between-subject variability *V*_b_ and within-subject variability *V*_W_. Small circles indicated the mean ICC of all the vertices. The values of ICC are outlined with an interval of 0.1 in the plane. b) Mean between-subject variability of all vertices within the seven canonical large-scale brain networks are plotted for both 3T (solid line) and 7T (dash line) magnet while the face colors of these circles indicate the level of *V*_b_: Visual (visual), SomMot (sensory motor), VentAttn (ventral attention), DorsAttn (dorsal attention), Control (frontal parietal control network), Limbic (limbic) and Default (default). c) Mean within-subject variability of all vertices within the seven canonical large-scale brain networks are plotted for both 3T and 7T magnet while the face colors of these circles indicate the level of *V*_W_.

## Discussion

Under 3T magnets, the test-retest reliability has been demonstrated to be limited for measurements of CSA with rfMRI, especially the common functional connectivity metrics (ICC< 0.4, see a recent review in [32]). In contrast, some CSA metrics (e.g., ALFF, ReHo, VMHC and ICA) have shown moderate to substantial reliability (ICC> 0.6, see systematic reviews in [59, 61]). This may reflect the methodological differences across these metrics in mapping individual differences, and such a methodological choice can be an important factor of driving fMRI measurement reliability (see recent arguments on task-based fMRI in [15, 16, 29]). While the definitions of these metrics are different, several studies have reported high correlations among the three metrics when applied to mapping individual differences (e.g., [1, 50, 52]). Here, we presented evidence that ultra-high field (UHF) 7T MRI improves the ability to detect cortical spontaneous activity (CSA), thus exhibiting a higher test-retest reliability for mapping individual differences as well as distinctions among the metrics compared to 3T MRI.

We employed a spatial standardization method, namely the z-value transformation, to map individual differences in CSAs and to ensure the maps comparable across magnets and metrics. This method, which has been widely used for structural MRI image intensity normalization [34, 38, 39], decomposes an individual raw map of CSA into two components. The first component includes two global CSA measurements, i.e., mean and SD, of all the vertices across the cortex. The second component is a vertex-wise map of z-values, which is a spatially relative measurement of the CSAs normalized by the mean and SD. Our findings revealed that the UHF 7T magnet enhanced the signal-to-noise ratio of measuring the global CSA compared to the 3T magnet for all the different spatial scales. This is consistent with all the previous studies using fMRI under various conditions [22, 23, 28]. Beyond the global intensity increases of the CSA signals, different CSA metrics also demonstrated different patterns of changes on spatial distribution across the human cortical mantle. Specifically, the 7T magnet raised the spatial ranks of the lateral parietal areas in terms of their amplitude of CSA but maintained or lowered such ranks regarding the connectivity metrics of CSA. In contrast, the medial prefrontal cortex showed an inverse pattern of its spatial ranks. The direction of spatial ranks was different between the amplitude metric and the two connectivity metrics. Such metric-related differences were also observed regarding the test-retest reliability, which was lower for the amplitude metric under 7T magnet but higher for the two connectivity metrics although at the same level of measurement reliability. At both the global and the local vertex levels, it was clear that the reliability improvements were largely driven by the decreases in within-subject variability (i.e., the random errors) and the increases in between-subject variability (i.e., individual differences). This represents the first study of mapping changes in the individual differences in CSAs induced by the UHF 7T MRI comparing with the 3T MRI. Our findings also show a promising way of the z-value method for mapping individual differences, especially in multi-center studies (e.g., [11, 56, 58]).

The present work on the test-retest reliability of individual differences could inspire statistical considerations for experimental designs. As indicated by [61], given a statistical power of 80%, for a two-sample t-tests (widely used in case-control studies), a 10% increment of the measurement reliability (e.g., from 0.5 to 0.6 or 0.6 to 0.7) can save 50 samples when detecting a 0.3 experimental effect size. This is the case where the UHF 7T MRI offers such advantages over the 3T MRI, as we have demonstrated. In contrast, with the same number of samples, the use of the UHF 7T MRI can provide greater power to detect the effect size, and this has been demonstrated in previous studies of both functional activation [45] and effective connectivity experiments [43]. Beyond the above-mentioned points in theory, three aspects should be noted for measurement reliability in practice: the measurement target, measurement tool and measurement metric. When designing an experiment, a choice of the reliable measurements should be the first priority rather than simply a large sample size [61]. We conclude that the UHF 7T and 3T MRI can serve as robust tools for detecting highly reliable individual differences in human brain CSAs with the multi-scale metrics, while the UHF 7T MRI can even better detect connectivity metrics than 3T MRI.

## Acknowledgments

This work was supported by the National Basic Science Data Center ‘Chinese Data-sharing Warehouse for In-vivo Imaging Brain’ (NBSDC-DB-15). The neuroimaging data were provided by the HCP WU-Minn Consortium, which is funded by the 16 NIH institutes and centers that support the NIH Blueprint for Neuroscience Research 1U54MH091657 (PIs: David Van Essen and Kamil Ugurbil), the McDonnell Center for Systems Neuroscience at Washington University. We would like to thank Dr. Neda Jahanshad for her comments, suggestions, and for proofing the manuscript for English language.

## Author Contribution

Xiu-Xia Xing: Conceptualization, Formal analysis, Methodology, Writing - Original draft preparation. Chao Jiang: Data curation, Writing - Reviewing and editing, Visualization. Xiao Gao: Writing - Reviewing and editing. Yin-Shan Wang: Data curation, Writing - Reviewing and editing. Xi-Nian Zuo: Conceptualization, Writing - Reviewing and editing, Resources, Supervision, Project administration, Funding acquisition.

## Supporting Information

We provide all the supplementary figures here.

**Figure S1.**
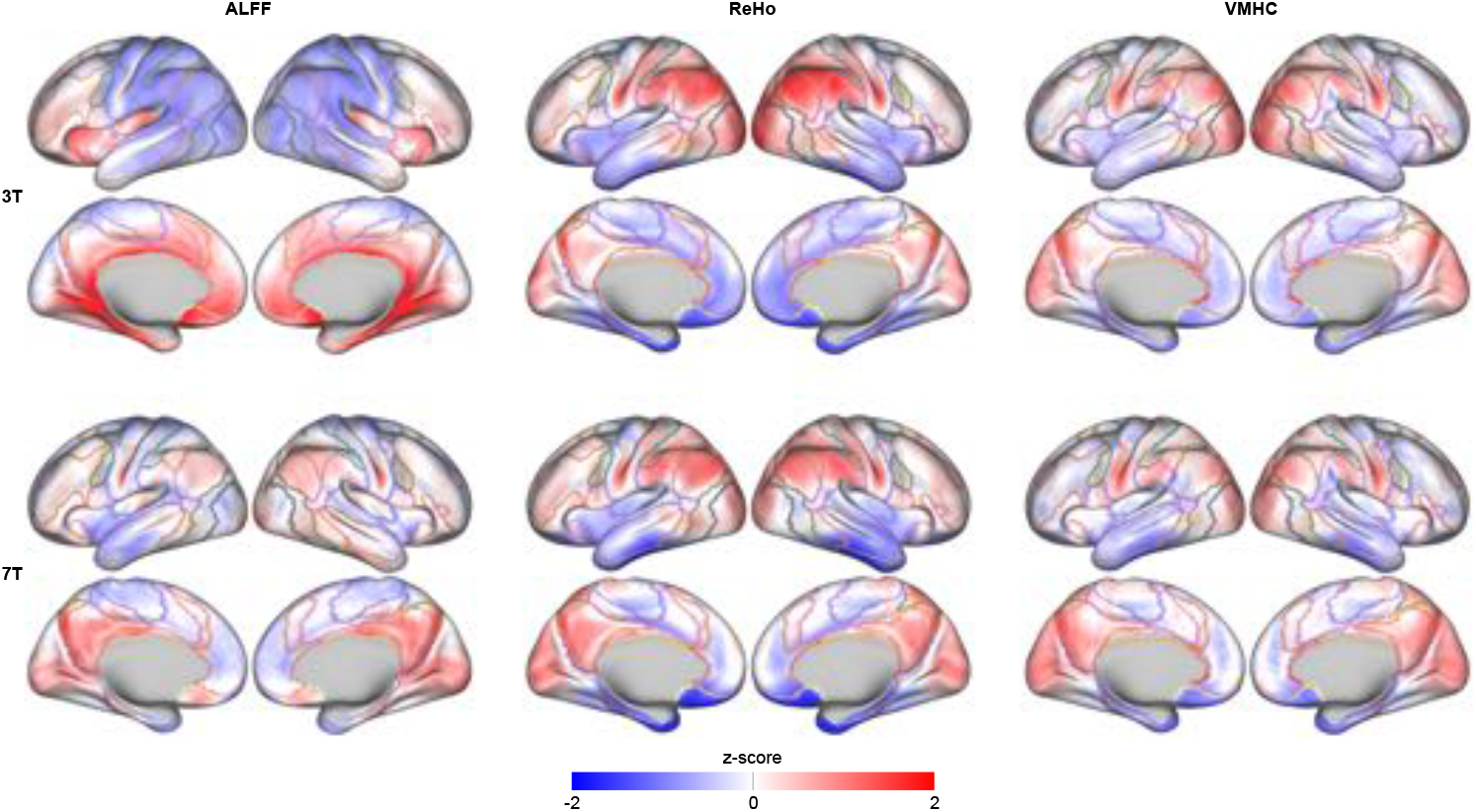
Vertex-wise mapping on mean z-values of the three CSA metrics.

**Figure S2.**
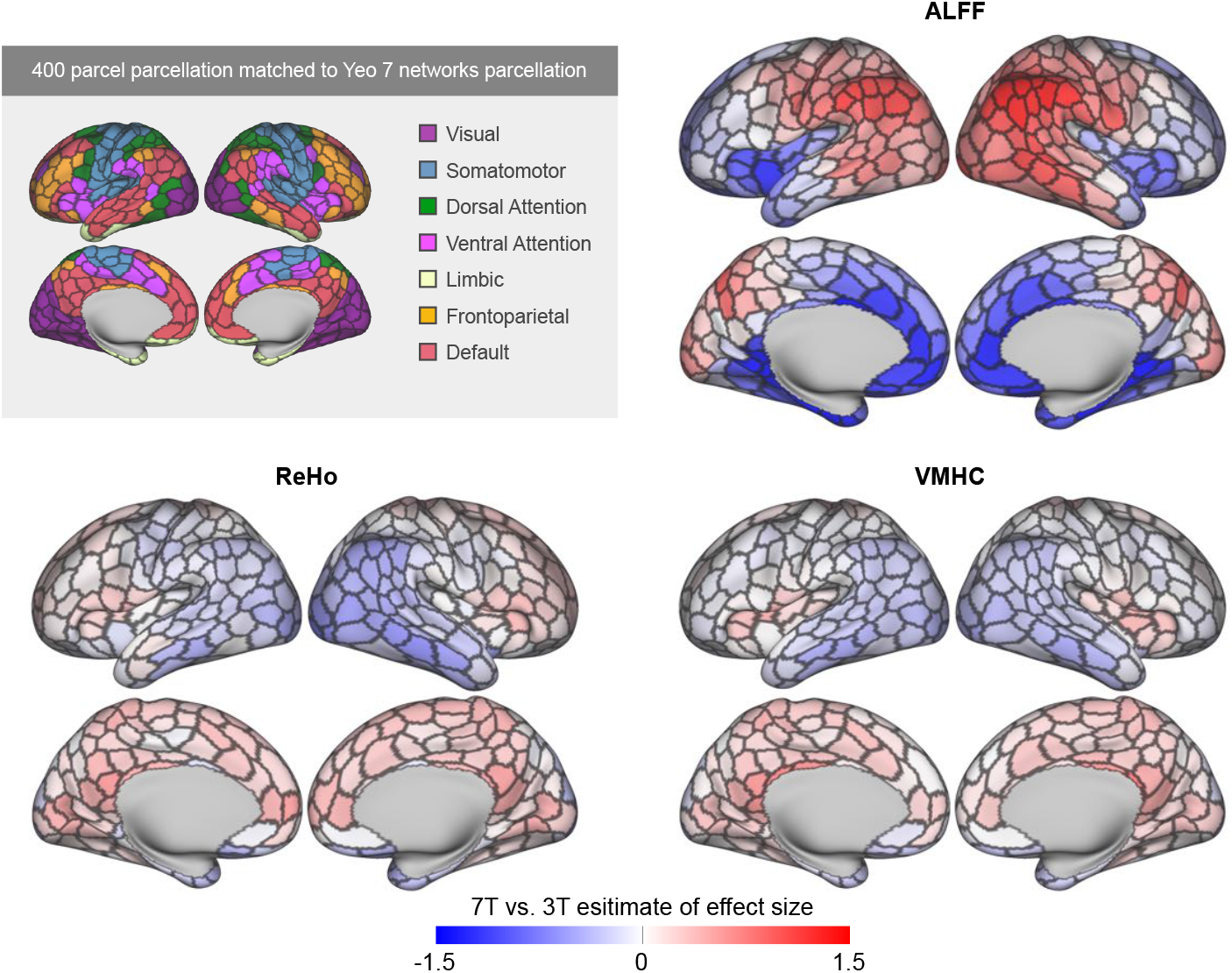
Parcel-wise mapping on differences in z-values of the three CSA metrics between 7T and 3T magnet.

**Figure S3.**
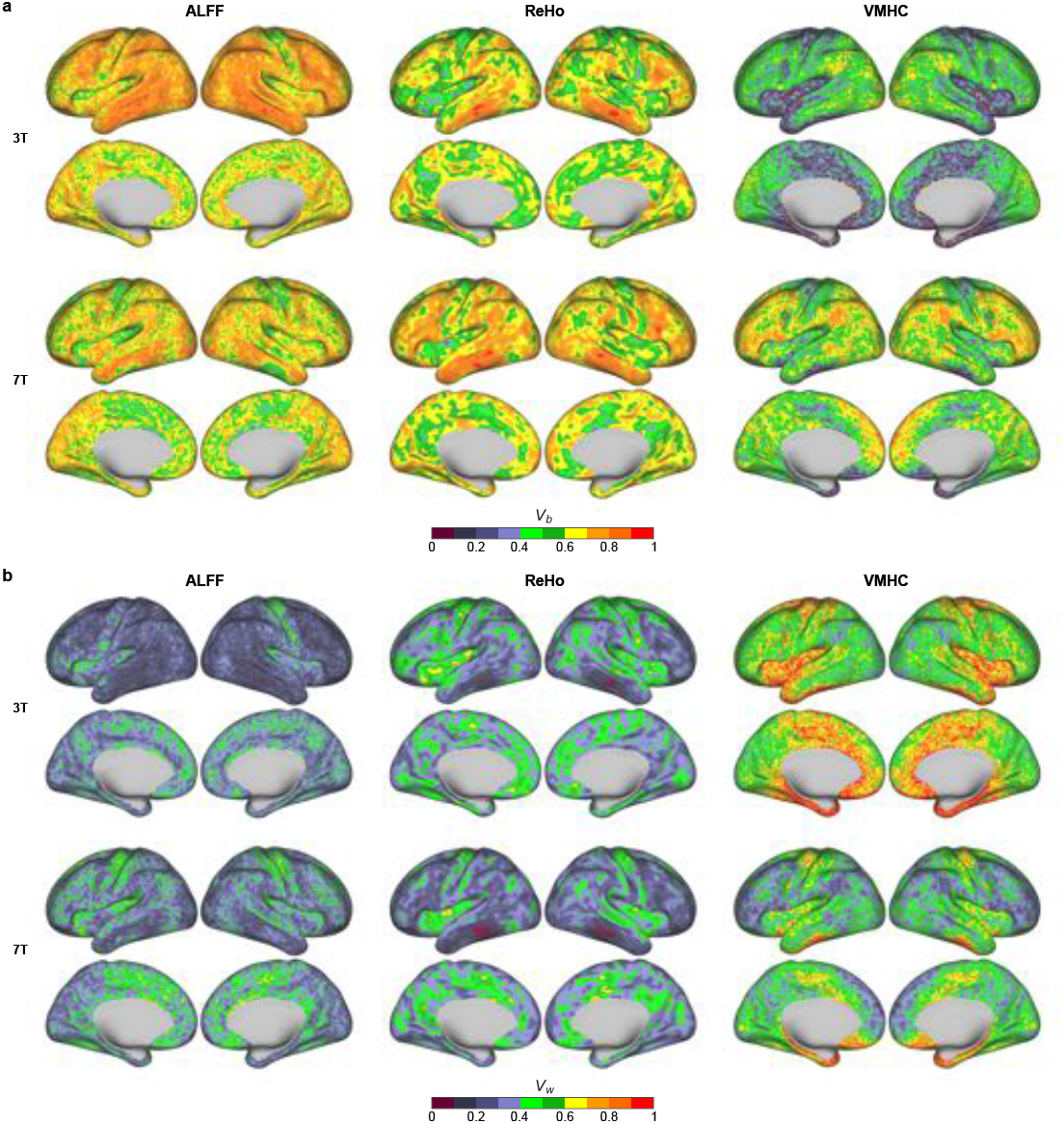
Vertex-wise mapping on individual differences in z-values of the three CSA metrics under both 7T and 3T magnet.

**Figure S4.**
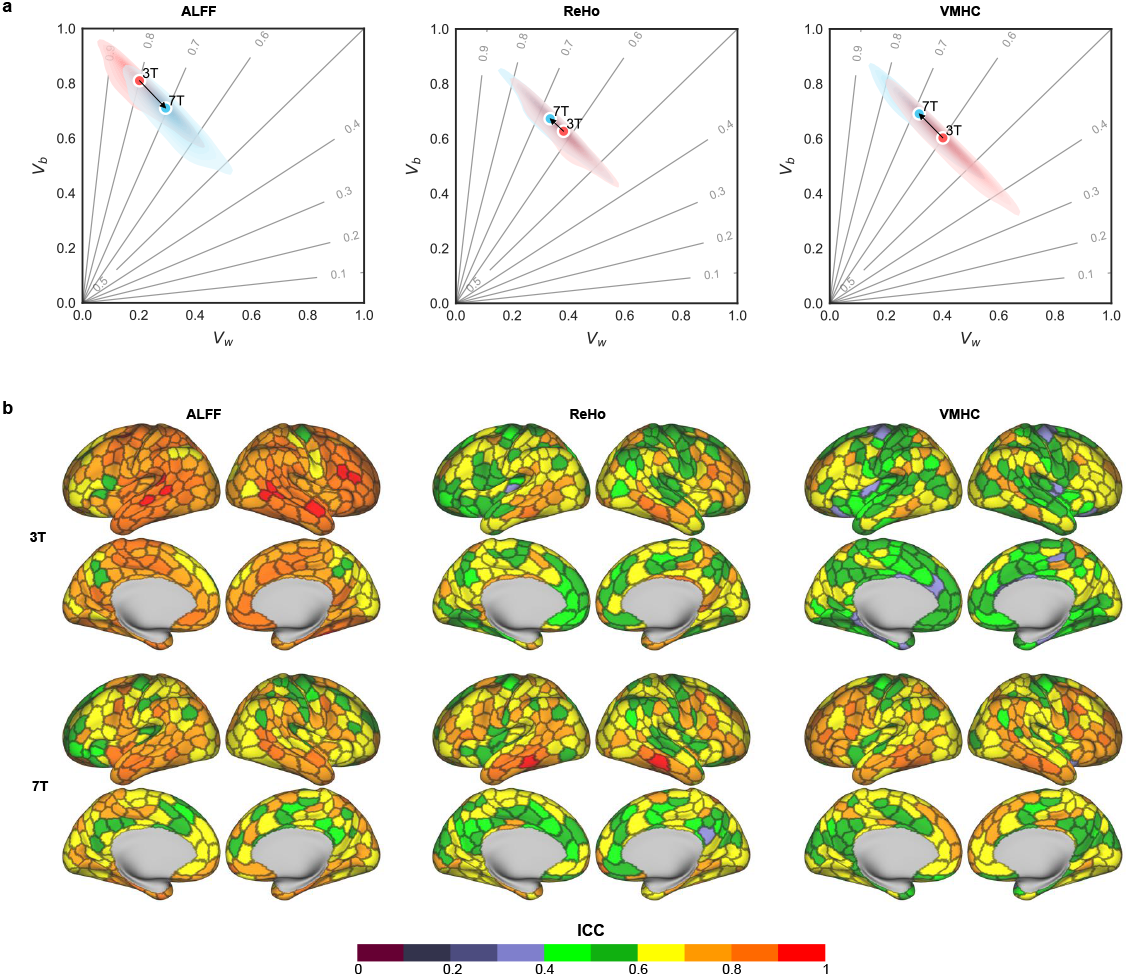
Parcel-wise mapping on test-retest reliability and individual variability of the three CSA metrics.

